# Cytoprotective roles of “E3 ubiquitin ligases-NF-κB-autophagy” axis in Pacific oysters *Crassostrea gigas* exposed to phenanthrene

**DOI:** 10.1101/2025.06.15.659737

**Authors:** Ziyi Guo, Zengguang Zhu, Hongyue Ma, Houqiang Du, Rong Tan, Weiwei Wang, Shaoguo Ru, Pengfei Cui

## Abstract

Phenanthrene (PHE), as one of the most frequently found polycyclic aromatic hydrocarbons can induce immunotoxicity, oxidative stress, and endocrine disruption in marine organisms. However, whether autophagy can be induced by PHE and the regulatory mechanism and cytoprotective roles of autophagy under PHE stress condition have not been unveiled. Our data first unveil a “E3 ubiquitin ligases-NF-κB-autophagy” axis, which play cytoprotective roles in Pacific oysters *Crassostrea gigas* exposed to PHE. The results of confocal laser scanning microscope, flow cytometry and transmission electron microscope confirmed that PHE could induce autophagy in the haemocytes of Pacific oysters, and the presence of autophagosomes was also confirmed. The proteomics results showed that the expression of the E3 ubiquitin ligase HUWE1, TRIM36, and autophagy-related protein 7 (ATG7) were significantly upregulated. The expression of genes of the “axis” were significantly upregulated, and the expression of genes of autophagy was downregulated after the inhibition of the NF-κB, indicating that the expression of the “axis”-related genes can be stimulated by PHE, and thus autophagy is activated. The upregulation of the expression of “axis”-related genes in mouse macrophages, further demonstrating the existence of the “axis” proposed by this study and the “axis” can be activated by PHE. Incorporating with changes of cell number, apoptosis rate, phagocytic capacity, and ROS levels of lymphocytes, we demonstrated that autophagy plays a cytoprotective role in cellular defence against PHE. This study proposed a novel pathway and supplied a comprehensive understanding of the protective role of autophagy in Pacific oysters to cope with pollutants.

## Introduction

Polycyclic aromatic hydrocarbons (PAHs) have significant effects on aquatic ecosystems and human health, and concerningly, are frequently found in marine environments. The United States Environmental Protection Agency (USEPA) has identified 16 PAHs, including phenanthrene (PHE), that require priority risk assessments and management, as they are classified as persistent organic pollutants (POPs) with highly toxic, carcinogenic, and mutagenic effects. PHE is one of the most frequently found PAHs in the environment, with concentrations of 2.44–26.1 μg/L (Hu et al., 2013; Liu et al., 2013). PHE is also readily bioaccumulated in organisms and biomagnified through the food web because of its lipophilic and persistent nature. The uptake of PHE can result in immunotoxicity, oxidative stress, and endocrine disruption in marine organisms) (Honda et al., 2021; Zhang et al., 2020). Among these creatures, bivalves have been shown to have a higher bioaccumulation and tolerance capacity for PHE than others, owing to their antioxidant and biotransformation enzyme-mediated detoxification systems (Zhong et al., 2016; Kroemer et al., 2010). Autophagy in bivalves can play more essential roles in the defence mechanisms, since it regulates and incorporates with the abovementioned factors through different signalling pathways (Zhong et al., 2016; Kroemer et al., 2010). Picot et al. (2019) have demonstrated that NH_4_Cl (one of the autophagy inducers) could cause autophagy in the blood lymphocytes of the Pacific oyster, *Crassostrea gigas*. Alexander et al. (2019) and Picot et al. (2020) identified thirty-five autophagy-related proteins in *C. gigas*. However, few studies have analysed the regulation pathways of autophagy in bivalves in relation to POPs, as well as its cytoprotective roles.

Autophagy regulation pathways in eukaryotes, including *C. gigas*, have been demonstrated to be similar to those in humans, including the AMP-activated protein kinase (AMPK) and Beclin 1/ class III phosphatidylinositol 3-kinase (PI3K) complex-related signalling pathways (Picot et al., 2019; Yang and Klionsky, 2010). Additionally, a recent approach has demonstrated that the NF-κB signalling pathway also maintains homeostasis and favours tissue repair by inducing autophagy, as it restrains its own inflammation-promoting activity and orchestrates a self-limiting host response (Zhong et al., 2016). The key proteins in the NF-κB pathway, such as IκB kinase (IKK) and the NF-κB inhibitor (IκB), are regulated by the ubiquitin-proteasome system (UPS). For example, E3 ubiquitin ligases TRAF6 and HOIP in the UPS can activate the IKK and NF-κB pathway, and ubiquitination mediated by the UPS causes the phosphorylation and degradation of IκB. This evidence demonstrates that the NF-κB signalling pathway can be activated by the UPS (Ikeda et al., 2011; Zemirli et al., 2014; Lechtenberg et al., 2016). Nevertheless, previous studies have mainly focused on elucidating the regulatory relationship between the NF-κB signalling pathway and autophagy and between the UPS and NF-κB signalling pathway. The relationships among the UPS, NF-κB signalling pathway, and autophagy, however, have not been revealed. The UPS and the autophagy-lysosomal pathway (ALP) are both major protein degradation pathways in eukaryotic cells, which are essential for maintaining homeostasis. The NF-κB pathway plays a central role in the regulation of cell survival and death by controlling immune responses and cell proliferation and development (Lechtenberg et al., 2016). Therefore, we hypothesised that the novel UPS-NF-κB-autophagy “axis” in bivalves play more essential role in protective and detoxification systems when exposed to POPs.

To demonstrate the hypothesis, we investigated the occurrence of autophagy in haemocytes of *C. gigas* after exposure to different concentrations of PHE using confocal fluorescent microscopy, flow cytometry, and transmission electron microscopy. Proteomics and real-time quantitative PCR were used to demonstrate the regulation pathways of autophagy in haemocytes of Pacific oysters exposed to PHE. The cell numbers, apoptotic rate, phagocytic rate, and phagocytic index of haemocytes were also measured to evaluate the self-protective role of autophagy in the haemocytes of Pacific oysters in response to PHE. The results have demonstrated that the UPS-NF-κB pathway-autophagy “axis” play key roles in the self-protection and detoxification in response to PHE toxicity, which provided a novel mechanism for the tolerance mechanism for organisms such as Pacific oysters to cope with pollutants.

## Results

### PHE Accumulates in Pacific Oyster Tissues

PHE was detected in all examined tissues of *C. gigas*, and its accumulation increased with higher exposure concentrations and longer durations (Fig. 1 and Supplementary Data S3). In circulating hemocytes (blood lymphocytes), PHE levels rose from about 0.9 ± 0.2 μg/L at 24 h with a low exposure (10 μg/L) to approximately 90 ± 12 μg/L after 7 days at the highest exposure (250 μg/L). This dose- and time-dependent uptake indicates that oyster hemocytes have a strong capacity to bioaccumulate PHE.

**Figure 1.**
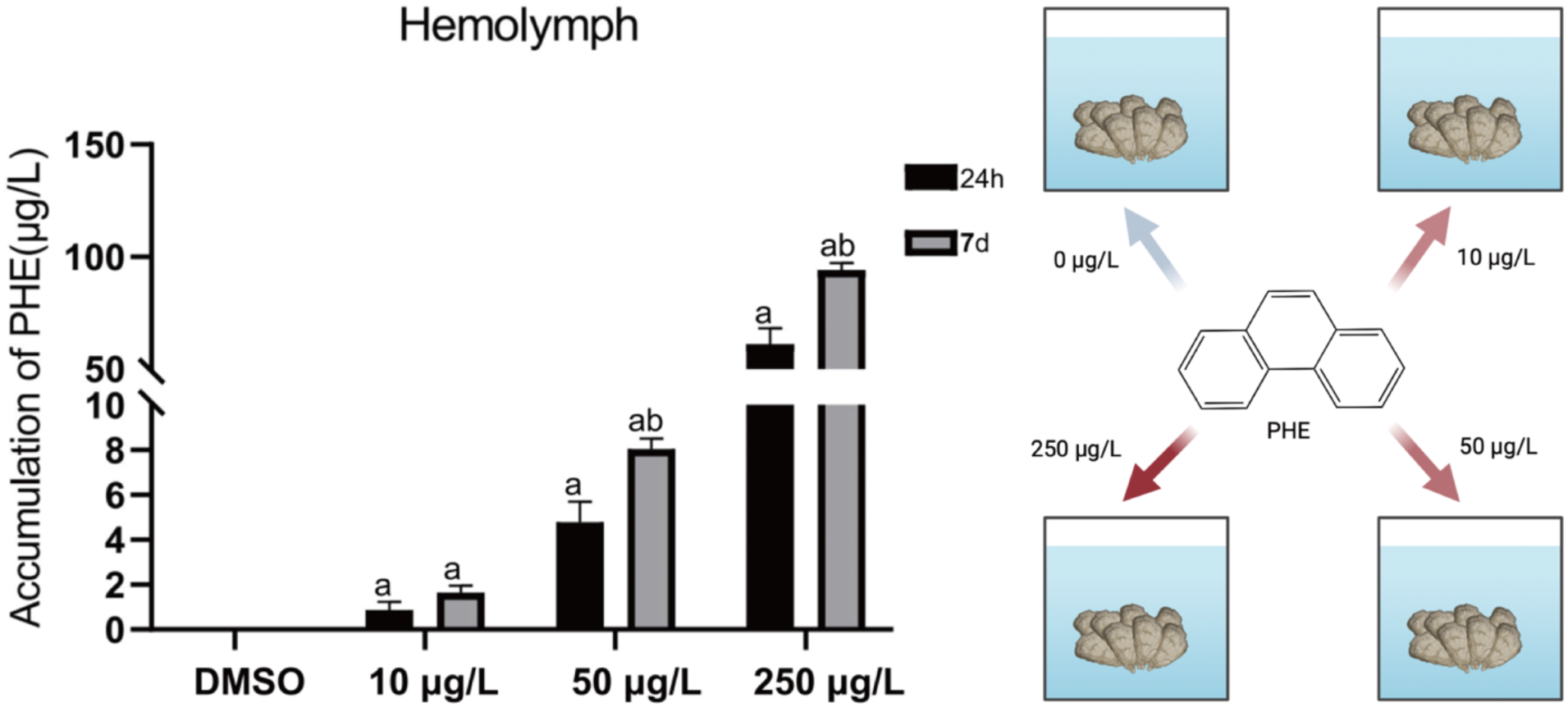
PHE concentrations in the haemolymph of *Crassostrea gigas.* ‘a’ represents a significant difference when compared to the control group, and ‘b’ represents a significant difference when compared to the 24 h exposure group (P < 0.05).

### PHE induces autophagy in oyster lymphocytes

#### CLSM approach to assess autophagic vesicles in the lymphocytes of Pacific oysters exposed to PHE

The presence of autophagic vesicles in the lymphocytes was evaluated under different conditions using CLSM by measuring the mean fluorescence intensity of 30 cells per condition. For the CLSM approach, the nuclei of the lymphocytes in the NH4Cl positive control group were stained with DAPI to emit blue fluorescence, autophagic vesicles showed green fluorescence, and numerous green-fluorescent vesicles appeared around the nuclei, which showed that numerous autophagic vesicles were produced in the lymphocytes. In the DMSO control group, only the nucleus emitted blue fluorescence, and there were no vesicles with green fluorescence inside the cells (Fig. 2A). Autophagic vesicles with green fluorescence were also observed around the nuclei of the lymphocytes in the 10, 50, and 250 μg/L PHE exposure groups, indicating the presence of autophagic vesicles in the lymphocytes.

**Figure 2.**
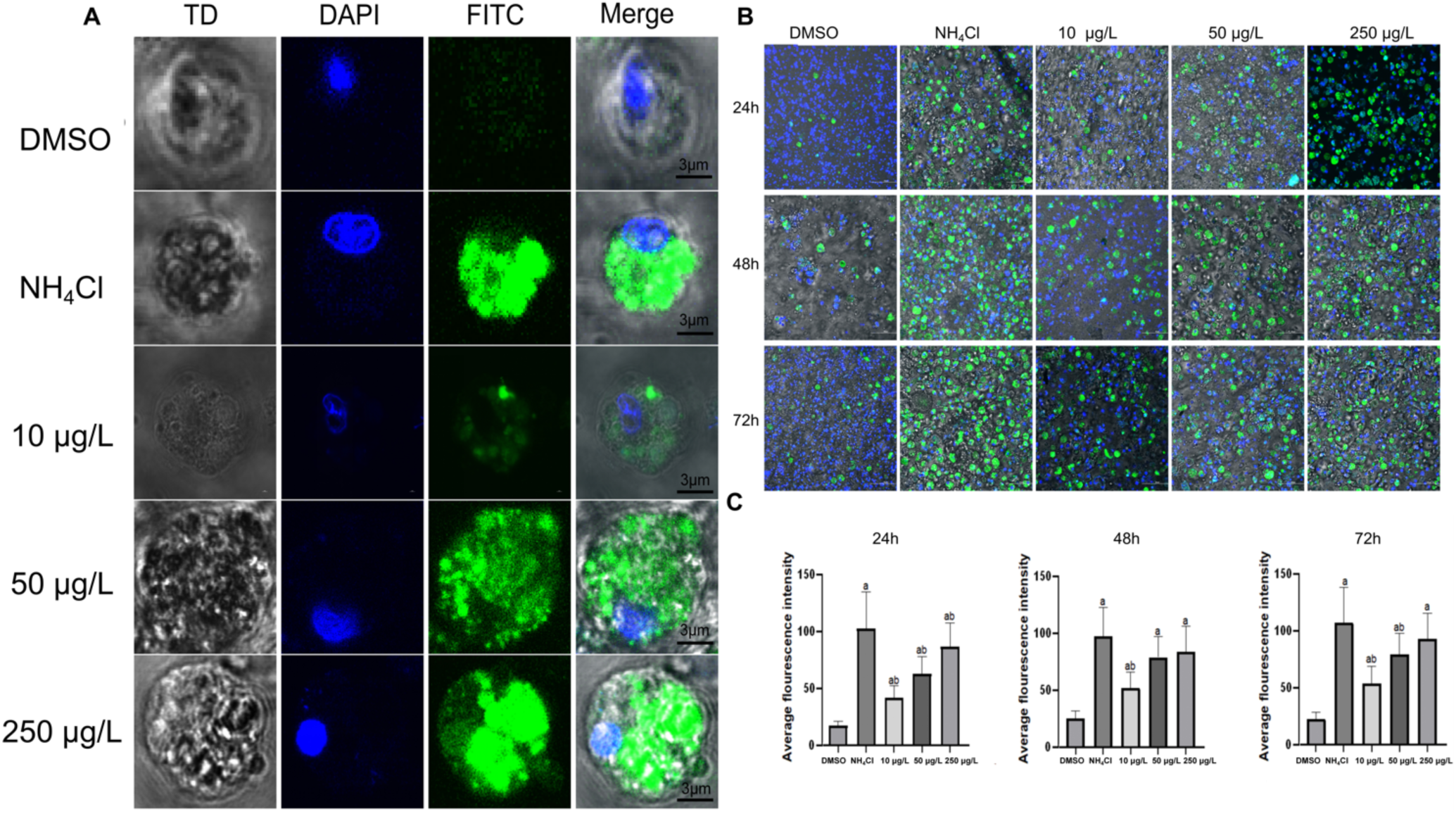
Observation and detection of the fluorescence intensity of haemolymph cells stained with Cyto ID®. (A) Pictures of the haemolymph under different experimental conditions (control, PHE, NH_4_Cl). The haemolymph was labelled with Cyto ID® (FITC) (column 3) and counterstained with 4′, 6-diamidino-2-phenylindole (DAPI) (column 2). Combined images of DAPI and FITC are presented in column 4. The edges of the blood lymphocytes can be observed in white light (TD) (column 1). Scale bar corresponds to 5 μm. (B) Confocal field of view of oyster haemolymphocytes stained using CytoID® after 24, 48, and 72 h exposures to the control, PHE, and NH_4_Cl. Scale bar corresponds to 50 μm (C) The mean fluorescence intensity of the haemolymph cells stained using CytoID® was calculated using Image J software, where ‘a’ represents a significant difference when compared with the control group, and ‘b’ represents a significant difference when compared with the NH_4_Cl-treated group, (P < 0.05).

After exposure for 24, 48, and 72 h, the mean fluorescence intensity in the lymphocytes of the NH_4_Cl positive control group was significantly increased when compared to that of the DMSO control group (P < 0.05). The green fluorescence intensity was also significantly increased in the lymphocytes exposed to PHE when compared to that in the DMSO control group (P < 0.05). The mean green fluorescence intensity of the 50 μg/L and 250 μg/L PHE exposure groups for 48 h and the 250 μg/L PHE exposure group for 72 h approached that of the NH_4_Cl positive control group (Fig. 2B and C).

#### Analysis of autophagic modulation in lymphocytes of C. gigas when exposed to PHE using flow cytometry

Two lymphocyte populations were identified: (1) negative lymphocytes, those not stained using Cyto ID®, and (2) positive lymphocytes, those that were stained using Cyto ID® (Fig. 3A). The percentage of lymphocytes stained using Cyto ID® differed according to the exposure conditions. These cell populations were further used to determine the autophagy activity of lymphocytes by calculating the ratios between the percentages of the cells with autophagosomes in the exposure and DMSO conditions (Fig. 3A). After exposure for 24, 48, and 72 h, the percentage of the cells containing fluorescence in the DMSO control group was 11 ± 2%, 8 ± 1%, and 7 ± 1%, respectively, with almost no fluorescence migration peaks. In the NH_4_Cl positive control group, however, the lymphocytes were found to be positive for staining using flow cytometry, and the percentage of cells containing fluorescence was 43 ± 3%, 38 ± 1.5%, and 34 ± 4%, respectively, with obvious fluorescence migration peaks. There was also significant fluorescence migration in the PHE exposure groups at 10, 50, and 250 μg/L, with the percentage of the fluorescence-containing cells at each exposure concentration for 24, 48, and 72 h being 26 ± 4%, 15 ± 2% and 19 ± 2%; 31 ± 2%, 25 ± 3%, and 24 ± 2%; and 35 ± 3%, 33 ± 1.5%, and 25 ± 4%, respectively (Fig. 3A). The results showed a significant increase (P < 0.05) in the proportion of fluorescence-containing cells in each exposure group when compared to that of the DMSO control group, with increasing exposure time and concentration (Fig. 3B). These flow cytometry data corroborate the CLSM findings, confirming that PHE exposure significantly induces autophagy in oyster hemocytes.

**Figure 3.**
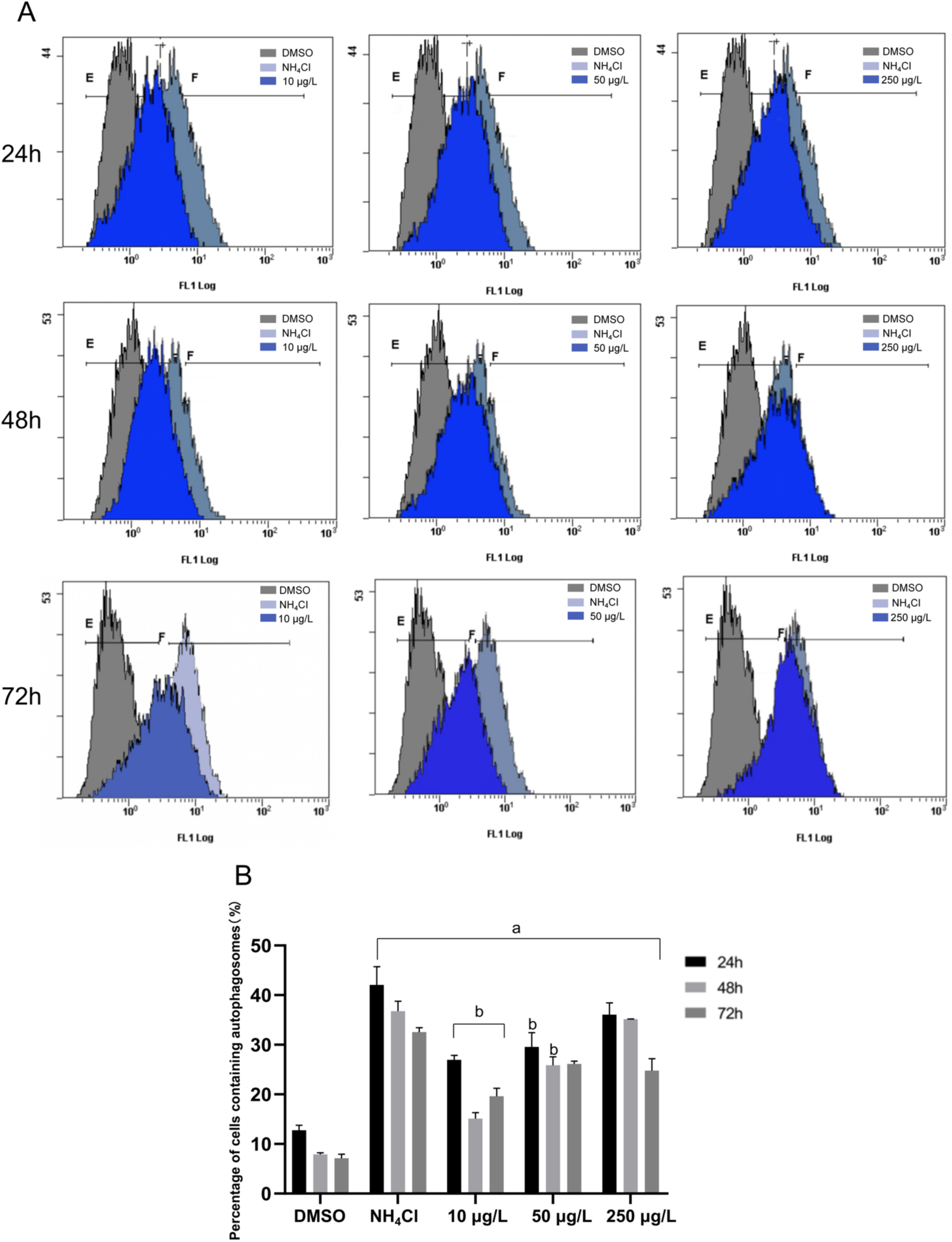
Detection and monitoring of autophagy in Pacific oyster blood lymphocytes using flow cytometry. (A) FL1 histograms of blood lymphocytes stained with Cyto ID® are superimposed. Normal state (control group) and autophagic vesicle staining (NH4Cl-treated group and 10 μg/L PHE-exposed group). The horizontal axis represents the fluorescence intensity, while the vertical axis represents the number of haemolymphocytes. Based on the Cyto ID® staining, two populations were defined: a negative cell population located on the left side of the histogram and a positive cell population located on the right side of the histogram. (B) 50 μg/L PHE-exposed group. (C) 250 μg/L PHE-exposed group. (D) Percentage of blood fine lymphocytes that are positive under the different test conditions (control, NH_4_Cl, 10 μg/L PHE, 50 μg/L PHE, and 250 μg/L PHE). Values are the means of six samples; error bars represent standard deviation. ‘a’ represents a significant difference when compared with the control, ‘b’ represents a significant difference when compared with the NH4Cl-treated group (P < 0.05).

#### Identification of ultrastructural modifications related to autophagy in the lymphocytes of C. gigas exposed to PHE using TEM

Ultrastructural examination by TEM further verified autophagy induction in PHE-treated hemocytes (Fig. 4). Control hemocytes exhibited normal morphology, with intact nuclei, non-swollen organelles, and only a few small vacuoles visible (Fig. 4A and F). In hemocytes exposed to PHE (72 h exposure, 10–250 μg/L), numerous double- or single-membrane autophagic vesicles were observed at all concentrations. At 10 and 50 μg/L PHE, cells showed an accumulation of autophagosomes accompanied by moderate vacuolization and condensation of nuclear chromatin (Fig. 4C and H, D and I). At 250 μg/L PHE, hemocytes contained abundant autophagosomes often found in proximity to or fused with lysosomes (indicating active autolysosome formation), and the nuclear membrane appeared partially disrupted in some cells (Fig. 4E and J). The NH₄Cl-treated positive control cells showed severe cytoplasmic vacuolation, enlarged endoplasmic reticulum, and numerous autophagic vesicles (Fig. 4B and G), closely resembling the autophagic ultrastructural changes reported in oyster hemocytes treated with NH₄Cl by Picot et al. (2019). These TEM results provide direct morphological evidence that PHE exposure induces the formation of autophagosomes and related autophagic structures in oyster hemocytes, consistent with activated autophagy.

**Figure 4.**
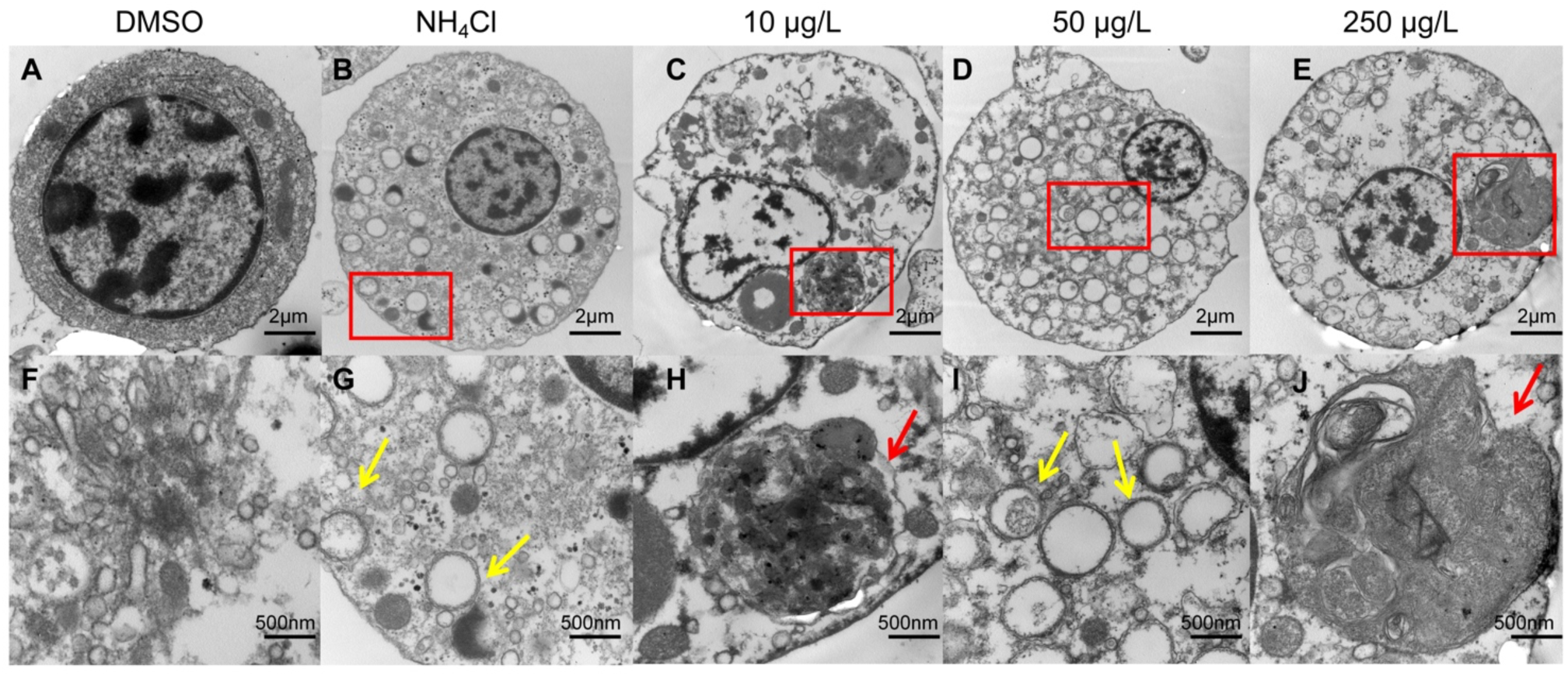
Electron microscopy images of Pacific oyster haemolymphocytes from the control, PHE-, and NH4Cl-treated groups for 24 h: (A) treated group, (B) NH_4_Cl-treated group, (C) 10 μg/L PHE-exposed group, (D) 50 μg/L PHE-exposed group, (E) 250 μg/L PHE-exposed group, (F) magnified images of haemolymphocytes from the control group, and haemolymph cells form the (G) autophagic vesicles in the 10 μg/L PHE exposure group and (H) autophagic vesicles in the 50 μg/L PHE exposure group, (I-J). Lysosomal structures in the haemolymph cells in the 250 μg/L PHE exposure group. The red arrows in the images point to single-membrane-structure autophagosomes, and the blue arrows point to double-membrane-structure autophagosomes. The A-E scale bar corresponds to 1 μm, and the F-J scale bar corresponds to 500 nm.

### Proteomic analysis reveals autophagy-related proteins and pathways in pacific oyster lymphocytes exposed to PHE

To gain insight into the molecular pathways involved in PHE-induced autophagy, we performed a proteomic comparison of hemocytes from control vs. PHE-exposed oysters. Out of 3,875 proteins detected, 106 were significantly differentially expressed in the 50 μg/L PHE group compared to control (54 upregulated and 52 downregulated, P < 0.05; Fig. S2). Notably, among the most upregulated proteins were two E3 ubiquitin ligases (HUWE1-like isoform X1 and TRIM36 isoform X1) and the autophagy-related enzyme ATG7 (ubiquitin-like modifier-activating enzyme ATG7), suggesting activation of the ubiquitin-proteasome system and autophagy machinery by PHE.

Gene Ontology (GO) enrichment analysis of the differential proteins identified 19 significantly enriched biological process categories (P < 0.05), chiefly related to membrane organization, transmembrane transport, catabolic processes, protein folding and transcriptional regulation (Fig. 5). Functions associated with the cytoskeleton were broadly downregulated, whereas those involving microtubule organization and depolymerization were upregulated in PHE-treated hemocytes. These enriched functions align with processes integral to autophagosome formation and trafficking such as cytoskeletal remodeling and membrane transport.

**Figure 5.**
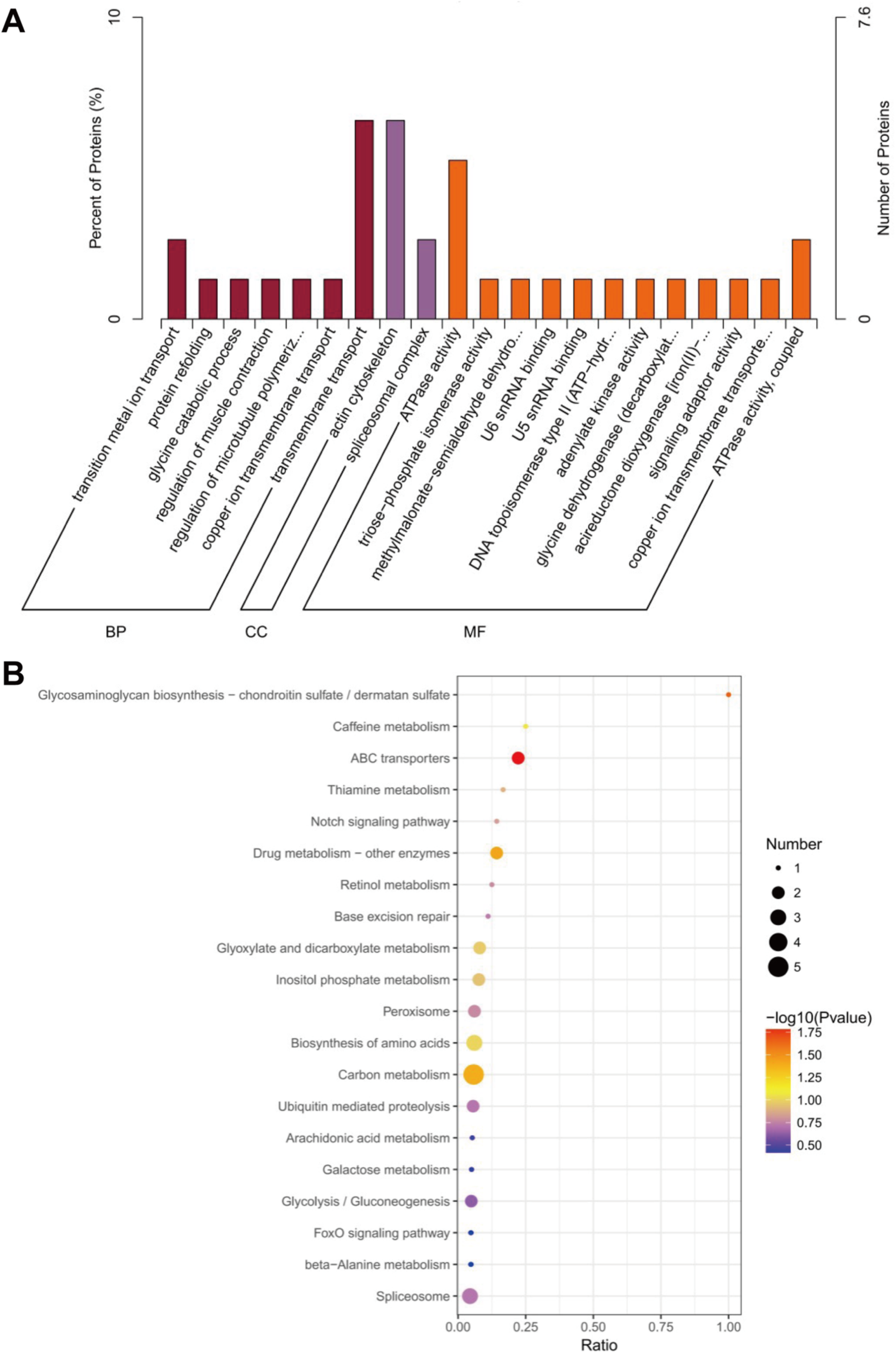
GO and KEGG Enrichment Analysis of PHE-Responsive Proteins. (A) GO classification analysis of the differentially expressed proteins (DEPs). Abscissa: GO functions; ordinate: the number and percentage of DEPs in each category. Red: Upregulated differential proteins in 50 μg/L PHE-exposed groups. Yellow: Downregulated differential proteins in group B. (B) KEGG classification analysis of the differentially expressed proteins (DEPs). The rich factor represents the ratio of the number of DEPs to the number of uniproteins in the pathway. The colour of the point represents the P value determined from the hypergeometric inspection. Red colour indicates a smaller value, and the size of the dot represents the number of differential proteins in the corresponding pathway, and the larger the dot, the more the number of differential proteins.

KEGG pathway analysis further showed five pathways significantly enriched among PHE-responsive proteins (Fig. 6): (1) ABC transporters (ABC transporters) and related proteins such as ATP-binding subunits; (2) Glycosaminoglycan biosynthesis-chondroitin sulphate/dermatan sulphate and related proteins such as carbohydrate sulfotransferases; (3) Drug metabolism-other enzymes and related proteins such as thiopurine-methyltransferases; (4) Carbon metabolism-related proteins such as 3-phosphoglyceraldehyde dehydrogenase, phosphotriose isomerase, peroxidase, and mitochondrial glycine dehydrogenase; (5) Caffeine metabolism-related proteins such as uricases. These results indicate that PHE exposure broadly alters cellular metabolism and transport processes in oyster hemocytes, alongside the upregulation of proteins directly implicated in the regulation of autophagy.

**Figure 6.**
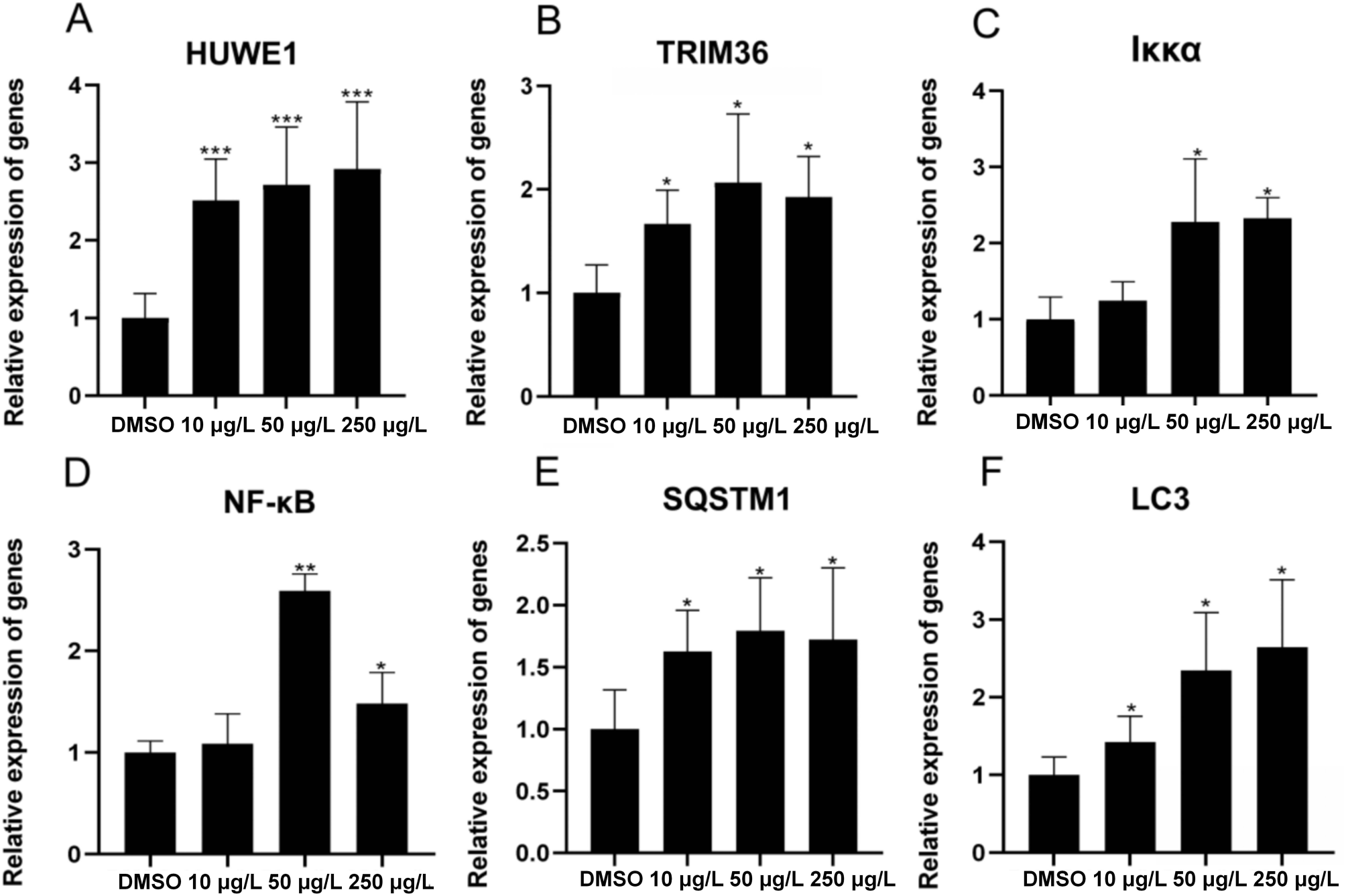
Results of real-time quantitative PCR assays of gene expression in Pacific oyster blood lymphocytes in the control group and each exposure group after 24 h. The symbols (***), (**) and (*) indicate P < 0.001, P < 0.01 and P < 0.05, respectively, compared with the DMSO group.

### Effects of PHE on the transcription of E3 ubiquitin ligases and NF-κB signalling pathway- and autophagy-related genes in Pacific oysters

#### PHE effects on the transcription of E3 ubiquitin ligase genes in Pacific oysters

Given the proteomic indications of an “E3 ubiquitin ligase–NF-κB–autophagy” regulatory axis, we next examined transcriptional changes in key genes of this axis in oyster hemocytes. Real-time quantitative PCR analyses showed that PHE exposure led to significant upregulation of genes encoding E3 ubiquitin ligases, NF-κB pathway components, and autophagy markers (Fig. 6). In particular, transcripts of genes encoding HUWE1 and TRIM36 (E3 ligases) were elevated at all tested PHE concentrations compared to control. At the highest dose (250 μg/L), *huwe1* mRNA was increased 2.78-fold (P < 0.01) and *trim36* approximately 2-fold (P < 0.05) relative to the DMSO control, confirming the proteomic finding that these ubiquitin ligases are induced by PHE.

#### PHE effects on the transcription of the NF-κB signaling pathway-related genes in Pacific oysters

The transcription of the NF-κB pathway-related genes in the lymphocytes was significantly increased in the 50 μg/L and 250 μg/L PHE exposure groups when compared with that of the DMSO control group (P < 0.05); however, there was no significant change in the 10 μg/L PHE exposure group, and the transcript levels of *Iκκα*, an upstream gene of NF-κB, were significantly increased in all exposure groups (P < 0.05; Fig. 6).

### PHE effects on the transcription of autophagy-related genes in Pacific oysters

Likewise, autophagy-related genes were upregulated: the key autophagosome component LC3 and the autophagy receptor p62/SQSTM1 coding genes both showed significantly increased transcription in PHE-treated hemocytes (P < 0.05 at all doses). Notably, *lc3* expression in the 250 μg/L group was 2.9-fold that of the control, indicating strong activation of autophagic machinery at the mRNA level (Fig. 6). These data support the notion that PHE triggers a concerted activation of the UPS–NF-κB– autophagy axis in oyster cells.

#### The transcription of E3 ubiquitin ligases and NF-κB signalling pathway- and autophagy-related genes in Pacific oysters after inhibiting the NF-κB signalling pathway

To test whether the induction of autophagy-related genes by PHE is dependent on NF-κB signaling, we inhibited the NF-κB pathway using NF-κB inhibitor CAPE and measured gene expression (Fig. 7). CAPE treatment alone significantly reduced basal NF-κB mRNA levels in hemocytes relative to uninhibited controls, confirming effective pathway suppression. When NF-κB was inhibited, PHE could no longer induce the full autophagic gene response. In PHE and CAPE co-treated hemocytes, the transcriptional upregulation of LC3 and p62 coding genes was abolished, their mRNA levels were significantly lower than in hemocytes exposed to PHE alone, and were comparable to baseline (Fig. 7E and F). In contrast, PHE-mediated induction of the E3 ubiquitin ligase genes was not significantly affected by NF-κB inhibition (Fig. 7C and D); HUWE1 and TRIM36 coding genes transcripts remained elevated even when NF-κB was blocked. These results indicate that PHE-induced autophagy gene upregulation (LC3, p62) requires an active NF-κB pathway, whereas the upregulation of the E3 ubiquitin ligases HUWE1 and TRIM36 coding genes by PHE occurs through an NF-κB-independent mechanism.

**Figure 7.**
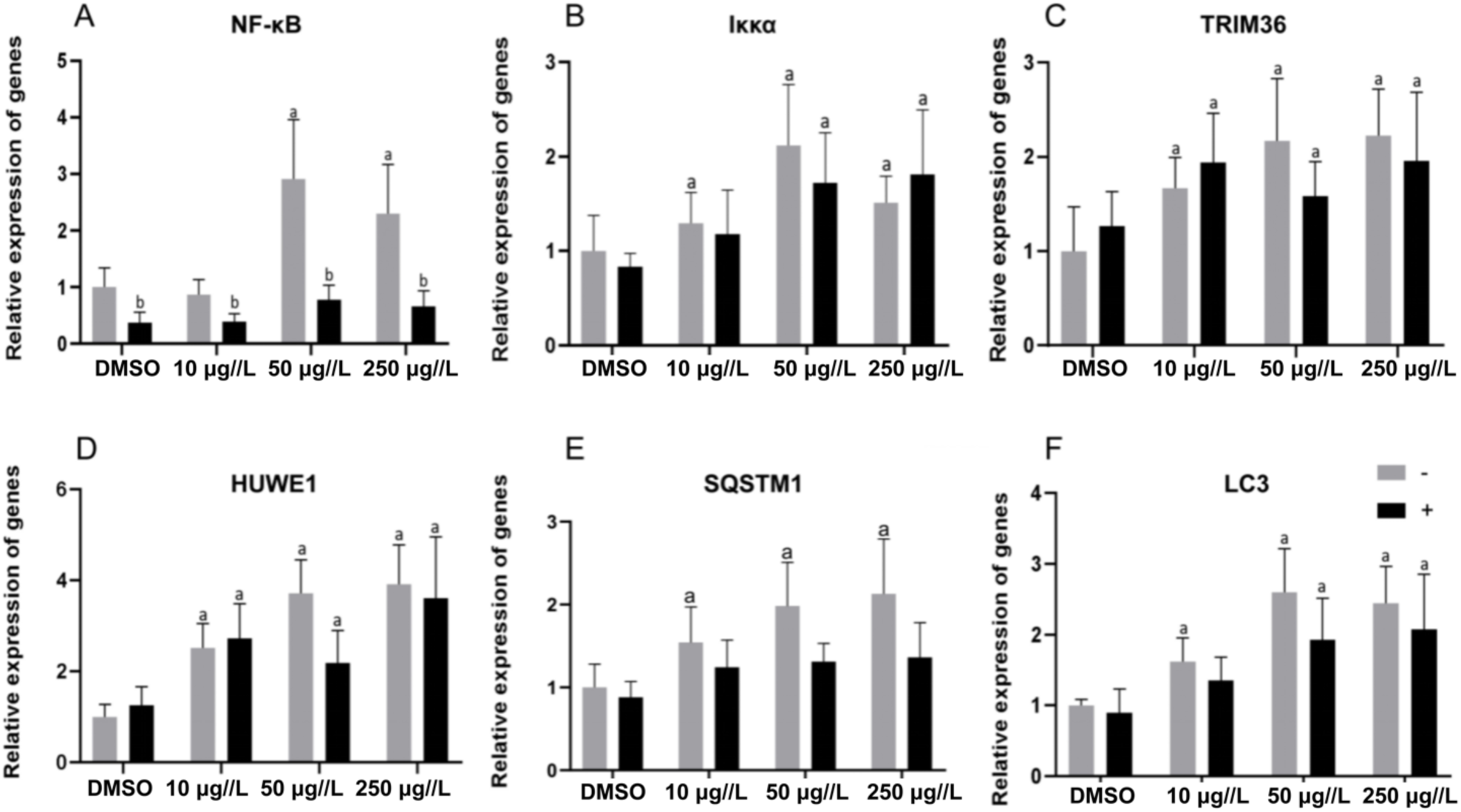
Results of real-time quantitative PCR assays of gene expression in Pacific oyster blood lymphocytes in control and exposed groups at 24 h after the NF-κB inhibitor (CAPE) was injected (black bars), when compared to that of the uninjected group (gray bars). ^a^Represents a significant difference from the control group (P < 0.05). ^b^Represents a significant difference from the uninjected group (P < 0.05).

### Effects of PHE on the transcription of E3 ubiquitin ligases and NF-κB signalling pathway- and autophagy-related genes in murine macrophages

We next asked whether the PHE-triggered “E3 ubiquitin ligase–NF-κB–autophagy” axis observed in oysters also operates in mammalian cells. Mouse macrophages (RAW264.7 cells) were exposed to PHE and examined for gene expression changes. Strikingly, PHE-exposed macrophages showed a significant upregulation of the same set of genes: the genes encoding the E3 ubiquitin ligases TRIM36 and HUWE1, NF-κB, and the upstream IKK, LC3, and p62/SQSTM1 were all transcribed at higher levels in PHE-treated cells than in DMSO-treated controls (P < 0.05 for each; Fig. 8). This conserved response suggests that PHE can activate the UPS–NF-κB–autophagy signaling axis across species, and it further supports the existence of this regulatory triad in the cellular defense against PHE toxicity.

**Figure 8.**
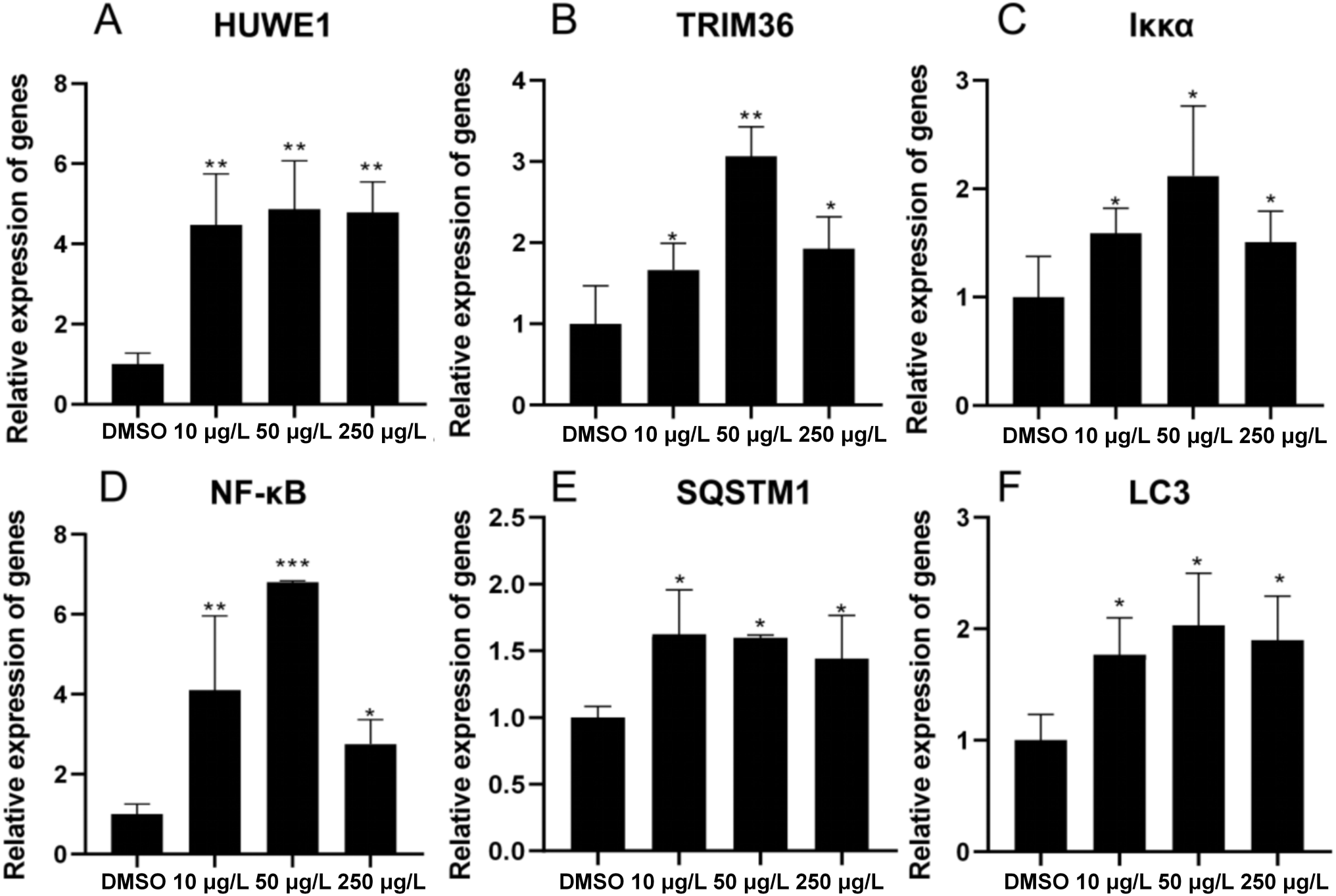
Real-time quantitative PCR results for genes in mouse macrophages in the control group and each exposure group after 24 h. The symbols (***), (**) and (*) indicate P < 0.001, P < 0.01 and P < 0.05, respectively, compared with the DMSO group.

### Effect of PHE on the total number of lymphocytes and their apoptosis rate, phagocytic capacity, and ROS levels in the Pacific oysters

To evaluate the impact of PHE on oyster immune cell viability and function, we measured hemocyte (lymphocyte) counts, apoptosis rates, phagocytic activity, and reactive oxygen species (ROS) levels after exposure (Fig. 9). After 24 h of PHE treatment, the total number of hemocytes was significantly higher in all PHE-exposed groups compared to the DMSO control (Fig. 9A). However, the magnitude of this increase was inversely related to PHE concentration – lower doses caused a larger rise in cell count – suggesting that higher PHE levels may begin to impair cell proliferation or survival. By 48 h, hemocyte counts in the PHE groups had declined to levels statistically indistinguishable from the control, indicating the initial boost was transient.

**Figure 9.**
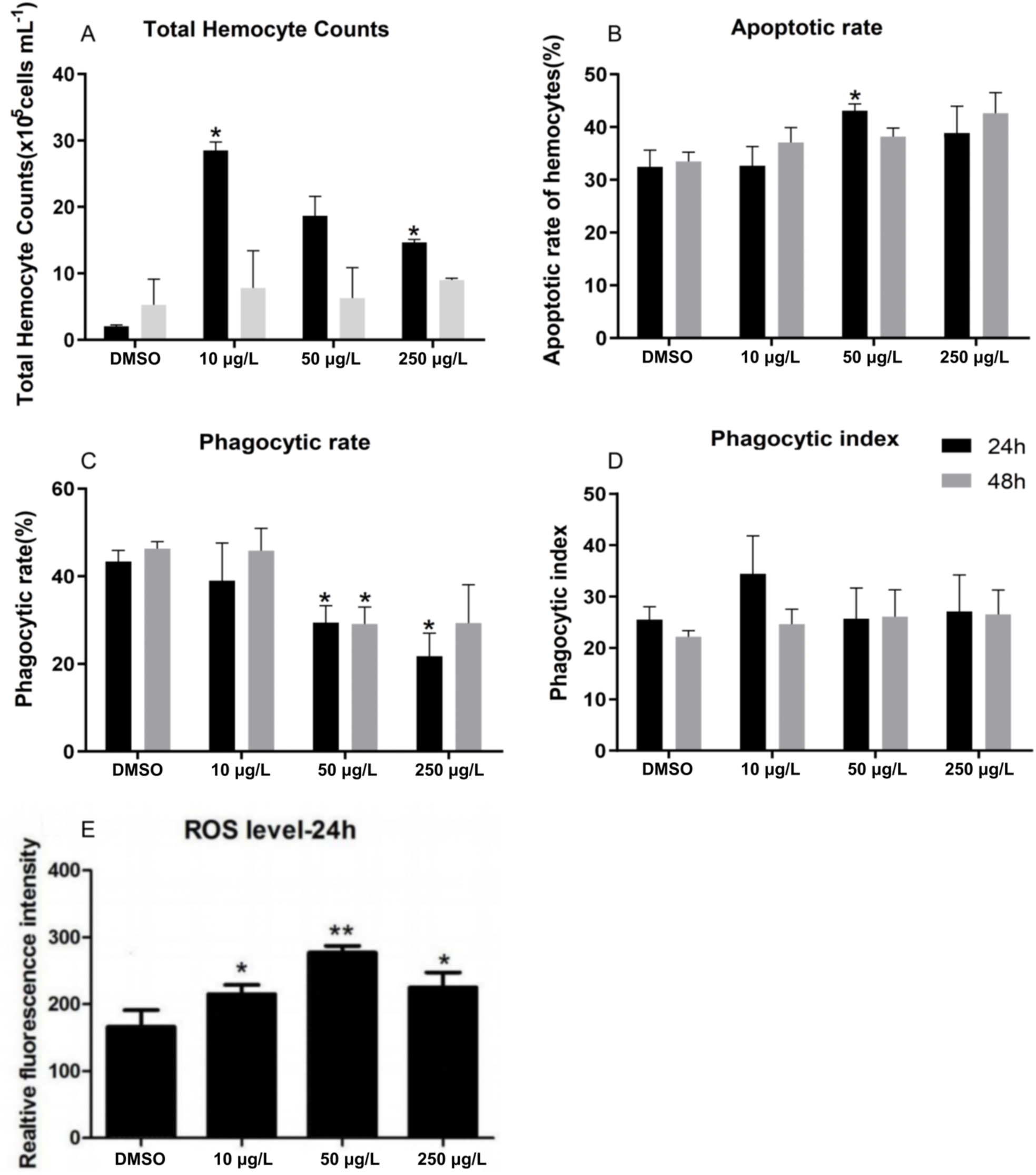
Effects of phenanthrene (PHE) on the total number of (A) Pacific oyster blood lymphocytes, (B) apoptosis rate, and (C, D) phagocytic capacity. (E) ROS values in Pacific oyster blood lymphocytes after 24 h of control (DMSO) and PHE exposure. Values are expressed as mean ± standard deviation (n = 3). The symbols (**) and (*) indicate P < 0.01 and P < 0.05, respectively, compared with the DMSO group.

PHE exposure had a relatively mild effect on hemocyte apoptosis. At 24 h, only the mid-dose PHE group (50 μg/L) showed a significant increase in the percentage of apoptotic cells compared to control, whereas no significant change was observed at 10 or 250 μg/L (Fig. 9B). By 48 h, apoptosis rates in all PHE-treated groups were similar to control levels, suggesting any pro-apoptotic effect of PHE was limited and transient.

PHE did, however, transiently suppress the phagocytic capacity of oyster hemocytes. After 24 h exposure, the phagocytic rate (the proportion of hemocytes engulfing fluorescent beads) was significantly reduced in the 50 and 250 μg/L PHE groups relative to the control (Fig. 9C and D). The 10 μg/L group showed no significant change in phagocytosis at 24 h. After 48 h, phagocytic activity partially recovered: the 50 μg/L group still exhibited a lower phagocytic rate than the control, but the 10 and 250 μg/L groups showed phagocytic rates comparable to control values. The phagocytic index (average number of particles ingested per phagocytic cell) was not significantly affected by PHE at either time point, indicating that for those cells that did carry out phagocytosis, their capacity on a per-cell basis remained similar to controls. Lastly, ROS production in hemocytes was strongly influenced by PHE. Intracellular ROS levels were significantly elevated in all PHE-treated groups after 24 h compared to the control (Fig. 9E). This elevation of ROS suggests that PHE exposure induces oxidative stress in oyster hemocytes. Elevated ROS, together with the observed changes in cell counts, apoptosis, and phagocytosis, underscores the immunotoxic and stress effects of PHE on oyster cells. However, taken in context with the activation of autophagy noted above, these results also hint that autophagy may be acting to counterbalance PHE’s cytotoxic effects, a possibility further addressed in the Discussion.

## Discussion

PHE is the smallest PAH with a bay- and K-region, and it also has highly reactive regions where the main carcinogenic species can be formed; furthermore, it is readily bioavailable, exhibiting the characteristic of bioaccumulation in aquatic organisms (Hannam et al., 2010; Ning et al., 2010). The results of this study indicate that the accumulated concentrations of PHE in the haemocytes of Pacific oysters could reach 0.9 ± 0.2 μg/L after exposure to PHE at the environmentally relevant concentration of 10 μg/L for 24 h, while the concentration reached 90 ± 12 μg/L after exposure to 250 μg/L of PHE for 7 d. These results indicate that the haemocytes of Pacific oysters have a strong capacity to accumulate PHE. The haemolymph is the open circulatory system of Pacific oysters, in which all tissues infiltrate, and it can act as a transmission medium for contaminants and corresponding metabolites (Pan et al., 2006). Numerous studies have demonstrated that PHE has toxic effects, such as immunotoxicity, causing embryonic abnormalities, and genotoxicity, in wildlife, including fish, benthic organisms, and marine vertebrates (Machado et al., 2014; Romero et al., 2018). Studies have demonstrated that PHE at concentrations of 0.2, 1.0, and 5.0 μg/L could cause adverse effects on reproduction and impair the development of juvenile zebrafish (Peng et al., 2019). In addition, PHE at concentrations of 1 μM induced cardiotoxic effects in zebrafish by impacting cardiomyocyte excitability (Peng et al., 2019). Furthermore, PHE has been shown to cause liver morphology deterioration and cellular damage in exposed zebrafish (Mai et al., 2019). Unlike the significant toxicity that zebrafish experience with PHE, bivalves have the notable ability to bioaccumulate and tolerate PHE and other pollutants. A previous study reported that PHE treatments did not produce any significant alterations to the levels of the antioxidants (CAT and GPx) and related enzymes (GR and G6PDH) in the gill or digestive gland and the expression of GST genes in the digestive gland of *C. brasiliana* when treated with both 100 μg/L and 1000 μg/L PHE (Lüchmann et al., 2014). These results indicate that oysters have strong self-protection and repair abilities when exposed to PHE.

Recent studies have shown that autophagy constitutes a major protective mechanism that allows cells to survive by eliminating potentially toxic substances and improving their adaptability in response to multiple stressors and helps defend organisms against pollutants and inflammatory and infectious parameters, playing a central role in protecting cells and maintaining homeostasis (Lroemer et al., 2010; Wang et al., 2018; Alexander et al., 2010; Yun et al., 2020). In this study, the CLSM observations showed that autophagosomes with green fluorescence were detected in the haemocytes of Pacific oysters exposed to PHE at different concentrations, and the number and fluorescence intensity of the autophagosomes showed a concentration-dependent effect with PHE; the results of the flow cytometry showed that the proportion of cells containing autophagosomes increased significantly with the increase in PHE concentration. Both the qualitative and quantitative data from the CLSM and flow cytometry confirmed that PHE could induce autophagy in the haemocytes of Pacific oysters. The presence of autophagosomes was confirmed using TEM, and autolysosomes and autophagosomes containing damaged organelles were also observed. Double- and single-membrane-bound vacuoles resembling autophagic structures were observed in all conditions, in which the autophagosomes had a bilayer structure, and the number of autophagosomes increased with the PHE concentrations. It is of note that the effects of inducing autophagy at 50 μg/L and 250 μg/L PHE were similar to those of NH4Cl. Therefore, we believe that autophagy is a newly identified self-protection mechanism in the haemocytes of Pacific oysters in response to PHE stress.

The NF-κB signalling pathway is an essential regulatory pathway of autophagy (Kretowski et al., 2016), and E3 ubiquitin ligases activate the NF-κB signalling pathway via the activation of IKK and the degradation of IκB (Zemirli et al., 2014). However, at present, there is no evidence available to elucidate the correlation among E3 ubiquitin ligases, the NF-κB signalling pathway, and autophagy. We have proposed that the E3 ubiquitin ligases-NF-κB pathway-autophagy “axis” can activate and regulate autophagy and plays a cytoprotective role in cells exposed to PHE. The proteomics results showed that the expression of the E3 ubiquitin ligase HUWE1 was significantly upregulated (P < 0.01), as was that of the TRIM36 and ATG7 in the family of autophagy-related proteins (P < 0.05). ATG7 is an autophagy-related protein that is crucial for the formation of autophagy coupling systems (Cawthon et al., 2018; Nakatogawa et al., 2009). These results indicate that the E3 ubiquitin ligases HUWE1 and TRIM36 promote autophagy. E3 ubiquitin ligases (E3s) are the most heterogeneous class of enzymes in the ubiquitination pathway (there are > 600 E3s in humans), as they mediate substrate specificity. Currently, E3 ubiquitin ligases can be classified into three main types, RING E3s, HECT E3s, and RBR E3s, depending on the presence of characteristic domains and the mechanism of ubiquitin transfer to the substrate protein. E3s can promote specific ubiquitination modifications of core proteins in relevant pathways according to different substrates in the stress response and plays an essential role in autophagy, immune signalling, oxidative stress, and other processes (Cheng et al., 2015; Morreale and Walden, 2016). The E3 ubiquitin ligase HUWE1, which was significantly expressed as per our proteomics results, belongs to the HECT E3 ubiquitin ligase family, while TRIM36 is a member of the RING E3 ubiquitin ligase family. Real-time quantitative PCR results showed that the expression levels of genes encoding E3 ubiquitin ligase, HUWE1 and TRIM36, IKK, and NF-κB, as well as autophagy receptors, LC3 and p62, in lymphocytes of Pacific oysters were significantly increased after PHE exposure, indicating that the expression of the axis (E3 ubiquitin ligases-NF-κB pathway-autophagy)-related genes can be stimulated by PHE, and thus autophagy is activated. Meanwhile, if the expression of related genes in the PHE-induced “E3 ubiquitin ligases-NF-κB pathway-autophagy” axis was inhibited by relevant inhibitors, autophagy would be down-regulated accordingly. In accordance with the above hypothesis, our results verified that the expression levels of genes encoding IKK and E3 ubiquitin ligases upstream of the NF-κB pathway were not significantly altered when the NF-κB signalling pathway was inhibited by the inhibitor, while the expression levels of autophagy-related genes were significantly downregulated, suggesting that autophagy was inhibited after the inhibition of the NF-κB signalling pathway. Autophagy is well-documented and conserved in most animals, and the macrophages of mice are a typical cell line for the study of autophagy (Huang et al., 2018). In this study, real-time quantitative PCR results of mouse macrophages showed that PHE could induce the upregulation of the expression of genes related to the “E3 ubiquitin ligases-NF-κB pathway-autophagy” axis, further demonstrating the existence of the autophagy regulatory pathway proposed by this study and that this axis can be activated by PHE.

Autophagy can degrade and recycle damaged organelles and cytoplasmic compounds through lysosomes, which can protect cells and contribute to the maintenance of cellular homeostasis under stress conditions (Zhong et al., 2016; Kroemer et al., 2010). Numerous autophagosomes and autolysosomes were observed via TEM in the haemocytes of Pacific oysters exposed to PHE, and many autophagosomes and autolysosomes containing damaged organelles were also observed, indicating that autophagy induced by PHE can recycle damaged organelles and play a role in protecting cells and maintaining cellular homeostasis. Additionally, autophagy can either regulate the phagocytosis of immune cells and other immune functions or remove intracellular ROS and damaged organelles to alleviate oxidative damage. Furthermore, autophagy can inhibit apoptosis through caspases, playing a cytoprotective role in stress response (Criollo et al., 2010; Marino et al., 2015). In this study, after exposure to PHE at different concentrations (10 μg/L, 50 μg/L, and 250 μg/L) for 24 h, the total number of blood lymphocytes (THC) increased significantly in the PHE exposure groups, the phagocytic index did not significantly change in the PHE exposure groups, the apoptosis rate did not significantly change in the 10 μg/L and 250 μg/L exposure groups, and the phagocytic rate decreased. After 48 h of exposure, the THC, phagocytic index, and apoptosis rate showed no significant changes, and the phagocytic rate gradually recovered. The intracellular ROS level was significantly increased after exposure to PHE at different concentrations (10 μg/L, 50 μg/L, 250 μg/L) for 24 h, and the ROS level in the 250 μg/L PHE exposure group was lower than that in the 50 μg/L PHE exposure group, indicating that the oxidative stress state of the cells was gradually restored. These results indicate that autophagy plays a cytoprotective role in the process of cellular defence against PHE, and the ability of haemolymph cells to phagocytose bacteria is not significantly affected, indicating that autophagy can maintain the immune defence function of haemolymph cells. Additionally, there was no significant change in the apoptosis rate of lymphocytes after PHE exposure, which may be related to the anti-apoptotic function of autophagy.

In this study, we proposed the “axis” of E3 ubiquitin ligases (HUWE1 and TRIM36)-NF-κB pathway (IKK and NF-κB in the NF-κB signalling pathway)-autophagy for the first time and identified that it plays a cytoprotective role in blood lymphocytes of the Pacific oysters *Crassostrea gigas* when exposed to PHE. PHE stimulated the expression of E3 ubiquitin ligases HUWE1 and TRIM36, and HUWE1 and TRIM36 sequentially activated IKK and NF-κB signalling pathways, followed by the recruitment of autophagy receptors LC3 and p62/SQSTM 1 to initiate autophagy (Figure 10). AMPK and PI3KC3-related signalling pathways are classical activation pathways of autophagy (Zhong et al., 2016). Studies have demonstrated that the NF-κB signalling pathway can regulate autophagy through AMPK and can induce the ubiquitination of Beclin1, a key protein in the PI3KC3 pathway, to promote autophagy (Hu et al., 2010). Therefore, the NF-κB signalling pathway in the “E3 ubiquitin ligases-NF-κB pathway-autophagy” axis proposed in this study could be a "bridge" to integrate with the classical regulatory pathway for autophagy (Kroemer et al., 2010). In addition, the E3 ubiquitin ligases in this axis can specifically ubiquitinate multifunctional signalling molecules such as IKK and NF-κB in the NF-κB signalling pathway in response to different forms of stress, coordinating and integrating autophagy and other stress responses (Zhong et al., 2016). Autophagy can be integrated with other cellular stress responses through the parallel stimulation of specific stress stimuli, the dual regulation of autophagy, and other stress responses by multifunctional stress signalling molecules and/or through the mutual control of autophagy and other stress responses. Thus, autophagy is an important mechanism that is a central component of the integrated stress response (Zhong et al., 2016; Kroemer et al., 2010). Therefore, the axis “E3 ubiquitin ligases-NF-κB pathway-autophagy” proposed by this study can coordinate and integrate autophagy and various stress responses during the response to sublethal stress. This integrated stress response is orchestrated through a multifaceted cellular program, by which cells undergo rapid changes to adapt their metabolism and protect themselves against potential damage. However, the mechanism of this "pathway/axis" in the integration of autophagy and stress responses in cells and in the self-protection against pollutants still needs further study.

**Figure 10.**
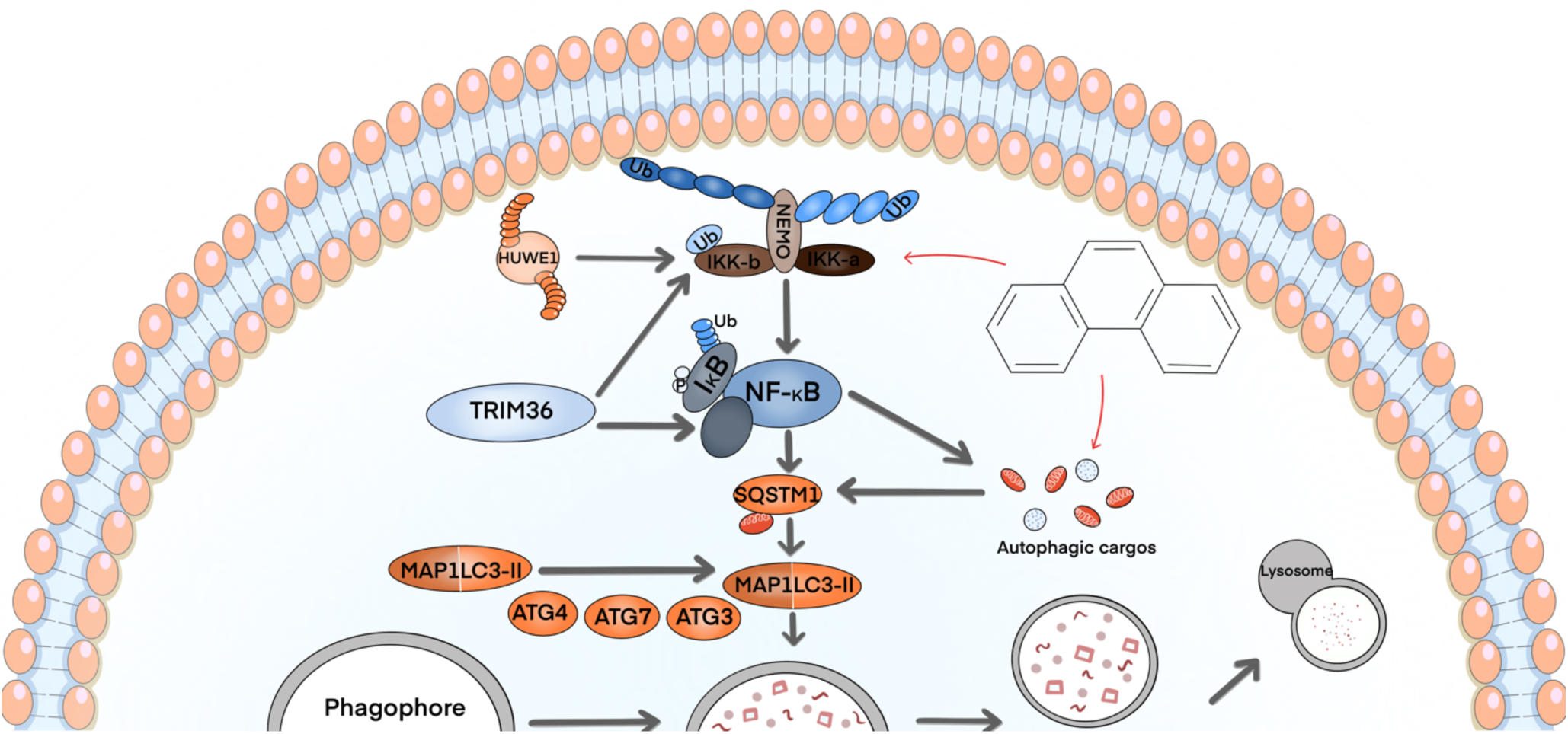
E3 ubiquitin ligases-NF-κB pathway-Autophagy axis/pathway in Pacific oysters *Crassostrea gigas* exposed to phenanthrene

## Materials and methods

### Experimental materials

Pacific oysters (*C. gigas*) were collected from a local oyster farm in Huangdao District, Qingdao, China (average shell length 7.3 ± 2 cm; average weight 90.2 ± 10 g). The oysters, after removing the organisms attached to their shells, were maintained in 50 L glass aquariums (40 oysters/tank), supplied with 20 L of 0.45 μm-filtered and continuously aerated seawater at 18 ± 2°C with a 12 h/12 h light/dark ratio. The water was also enriched in phytoplankton (*Spirulina platensis*) and changed daily prior to the experiments. The procedures for cultivating and using experimental organisms were approved by the Institutional Committee for the Protection and Use of Animals of the Ocean University of China. RAW264.7 cells were provided by the Stem Cell Bank of the Chinese Academy of Sciences. The cell culture medium was configured according to the “Mouse Leukaemia Virus-Induced Tumour Macrophages RAW 264.7 (SCSP-5036)”, and the culture medium contained 88% Dulbecco’s modified Eagle’s medium (DMEM) (Invitrogen, Carlsbad, CA, USA, 111960-044), 1% glutamax (Gibco, 35050-061), 1% sodium pyruvate (Gibco, 11360-070), and 10% foetal bovine serum (Gibco).

As PHE (CAS# 85-01-8, purity ≥ 99%, Shanghai Aladdin® Biochemical Science and Technology Co., Ltd., Shanghai, China) is insoluble in water, a solution (10 mg/mL) was first prepared by dissolving 100 mg of PHE in 10 mL of dimethyl sulfoxide (DMSO, Aldrich, Shanghai, China). The concentration of the PHE solution for the cell exposure experiments was 0.3 mg/mL, and it was stored at 25°C in brown glass bottles that were protected from the light, and PHE was renewed weekly.

### PHE exposure experiments and concentrations in Pacific oysters

#### PHE exposure experiments

Experiments were conducted using 50 L glass aquaria with 20 L of filtered seawater in each tank. The PHE solution (10 mg/mL) was added to the seawater to achieve final PHE concentrations of 10, 50, and 250 μg/L and a final maximum DMSO concentration of 0.025% (v/v). The PHE concentrations were added to the test media, and realistic environmental concentrations were selected based on previous bivalve research (Hu et al., 2013; Hannam et al., 2010; Lüchmann et al., 2012; Liu et al., 2013). Oysters were then randomly divided into glass exposure tanks (30 oysters in each tank), which were individually aerated and held for 24 h, 48 h, 72 h, and 7 d prior to the experiments. The DMSO control oysters were subjected to the same conditions as the exposed groups, except for the addition of 0.025% (v/v) DMSO only without PHE.

#### PHE concentrations in Pacific oyster tissues

The PHE concentrations were monitored over 24 h and 7 d to confirm that they were accurate. For each condition at each sampling time, 600 μL of haemolymph was extracted from the blood sinus of each oyster with a 1 mL sterile syringe. Three replicates of 600 μL haemocyte suspensions (approximately 1 × 10^5^ cells) were collected, and tissues from the gill, visceral mass, crustal membrane, and muscle were excised. The tissues were weighed and placed in 1.5 mL centrifuge tubes, then quick-frozen in liquid nitrogen, and stored at −80°C for subsequent experiments. The internal reference standard substances and 5 mL acetonitrile were mixed with the homogenates in each centrifuge tube. After vortexing and vibration mixing, ultrasonic solvent extraction was performed for 2 min. The homogenates were centrifuged at 2000 × g for 5 min, then the supernatants were collected, and the extraction procedure was repeated for the remaining residue. Anhydrous sodium sulphate was mixed with the supernatants, and they were then centrifuged at 2000 × g for 5 min, and then the supernatants were collected. After blow-drying at 40°C under nitrogen, the mixture was re-dissolved in dichloromethane and purified in a neutral alumina column which was activated with 3 mL of dichloromethane (1 g/6 mL) and then mixed with 5 mL of dichloromethane for elution. The re-dissolved solution and eluent were collected, after blow-drying with nitrogen, re-dissolved in 1 mL acetonitrile, and filtered through a 0.22-μm membrane. The PHE concentration was monitored using high-performance liquid chromatography (ChromasterUltra Rs, Hitachi, Japan) with a working wavelength of 254 nm and a mobile phase flow rate of 1 mL/min.

### Effect of PHE exposure on Pacific oysters’ lymphocytes autophagy

#### Exposure of Pacific oysters to PHE and NH_4_Cl

Experiments were conducted using 50 L glass aquaria with 20 L of filtered seawater in each tank. PHE solution (10 mg/mL) was added to the seawater to achieve final PHE concentrations of 10, 50, and 250 μg/L and a final maximum DMSO concentration of 0.025% (v/v). The NH_4_Cl (A801305; 99.8% purity, Macklin) positive exposure group was also set up, and the NH_4_Cl concentration was set at 0.053 g/L (Picot et al., 2019). Oysters were then randomly divided into glass exposure tanks (30 oysters per tank), which were individually aerated and held for 24, 48, and 72 h prior to the experiments. The exposure rearing conditions were the same as those used for pre-domestication.

#### Observation of autophagy in Pacific oyster lymphocytes after PHE exposure using confocal laser scanning microscopy (CLSM)

After 24, 48, and 72 h of exposure, six oysters were selected each from the DMSO control and the PHE and NH_4_Cl exposure groups. Haemolymph (800 μL) was extracted from the blood sinus of each oyster using a 1 mL sterile syringe and mixed into a 10 mL centrifuge tube, with the cell concentration reaching 1 × 10^6^ with the addition of phosphate-buffered saline (PBS). After centrifugation at 860 × g for 5 min, the supernatants were removed, and the pellets were pooled. The haemolymph cells were then incubated and stained according to the operating instructions of the Cyto ID® Autophagy Assay Kit (ENZO Life Sciences, ENZ-51,031-K200). After washing three times with PBS (pH7.4), the lymphocytes were observed under a fluorescence microscope at 60 × magnification using immersion oil (Nikon A1, Nikon, Japan). Images were extracted using NIS-Elements software (Nikon), and fluorescence intensity analysis was performed using Image J software (National Institutes of Health, USA). Histograms were generated, and statistical analyses were performed using GraphPad Prism version 5.0 (GraphPad Software, San Diego, CA, USA).

#### Assessing autophagy of Pacific oyster lymphocytes after PHE exposure using flow cytometry analysis Lymphocytes were suspended in 1 mL of buffer containing Cyto ID® Green Assay Reagent

(configured according to kit instructions), incubated for 30 min and stained at room temperature (21°C) for 90 min away from light and washed three times with PBS (pH7.4). The green fluorescence intensity of each group of cells was detected using flow cytometry (Beckman FC500-MPL, USA), and the lymphocytes containing autophagosomes emitted green fluorescence (FL1: 500-550 nm); three parallel samples were set up in each group, and the experiment was repeated three times. The parameters used for the analysis were based on the method described by Picot et al. (Picot et al., 2019). Two lymphocyte populations were identified using flow cytometry: (1) negative controls, lymphocytes not stained using Cyto ID®; and (2) staining group cells, lymphocytes stained using Cyto ID® and cells containing autophagic vesicles. The percentage of lymphocytes stained using Cyto ID® from the treatments with the different PHE concentrations were analysed to verify lymphocyte autophagy in the Pacific oysters when exposed to PHE.

#### Autophagy detection in the lymphocytes of Pacific oysters using transmission electron microscopy

Confocal microscopy results showed that the largest number of autophagic vesicles in the lymphocytes were identified after 72 h of PHE exposure. Twelve oysters were selected each from the DMSO control group and the PHE and NH_4_Cl exposure groups, after 72 h of exposure, and 800 μL of haemolymph was collected from the blood sinus with a 1 mL sterile syringe, mixed into a 10 mL centrifuge tube, with the cell concentration reaching 1 × 10^6^ with the addition of PBS. The prepared samples were fixed in 2.5% glutaraldehyde solution (Sigma-Aldrich, G5882) for 24 h at 4°C. Cells were subsequently washed three times with 0.4 M cacodylate buffer (Sigma-Aldrich, C0250) and post-fixed with a solution of 1% silver tetroxide (Sigma-Aldrich, 75,632) for 1 h at 4°C. The samples were then washed again in 0.4 M cacodylate buffer. After dehydration in successive baths of ethanol and gradual impregnation with propylene oxide, the samples were progressively impregnated and embedded in epoxy embedding medium (Sigma Aldrich, 45,345). After polymerisation at 60°C, the samples were cut into sections that wereof 1 μm thick for quality control and 80–85 nm thick for examination and then floated on a TEM grid, stained with uracil acetate/lead citrate (Lewis et al., 1977). The sections were observed using a transmission electron microscope (JEOL JSM-840) at 80 kV. The presence of ultrastructural modifications and autophagic structures in the haemocytes was examined using previously described criteria (Eskelinen and Kovács., 2011). Double-membrane-bound autophagic structures were identified based on the presence of a double membrane with a lumen between the two lipid bilayers, with contents resembling the cytoplasm around the structure in terms of density and composition.

### Proteomic detection of lymphocytes after PHE exposure

To investigate the regulatory mechanisms of PHE-induced autophagy in the lymphocytes of the Pacific oysters, proteomic analysis was performed. The results of the CLSM experiments showed that the largest number of autophagic vesicles were in the lymphocytes after 72 h of exposure to 50 μg/L PHE. Therefore, proteomic assays were performed using the DMSO group and the 50 μg/L PHE group with a 72-h exposure. Eight oysters were selected from each group, and 800 μL of haemolymph was collected from the blood sinus using a 1 mL sterile syringe. Each group was mixed into a 25 mL centrifuge tube with 800 μL of anticoagulant (PBS containing 10 mg/mL sodium heparin) and stored at 4°C. Three parallel samples were set for each group, and the methods of protein extraction, quality control, sequencing, and analysis were performed according to the iTRAQ method (Zhu et al., 2020) of Novozymes (Beijing, China).

### Real-time quantitative PCR analysis of expression of E3 ubiquitin ligases and NF-κB signalling pathway- and autophagy-related genes in Pacific oysters exposed to PHE and caffeic acid phenyl ether (CAPE)

#### Experimental methods for PHE and combined PHE and CAPE exposures to Pacific oysters

The PHE exposure method for the Pacific oysters was performed as described in Section “*PHE exposure experiments*”.

For the combined treatment, the PHE exposure was performed as described in Section “*PHE exposure experiments*”, followed by CAPE exposure using an injection method. Three PHE exposure concentration groups and a DMSO control group were set up, followed by the injection of 100 μL CAPE solution (configured with DMSO at 0.1 mg/mL) into the blood sinus of the Pacific oysters in each group with a 1 mL sterile syringe. The PHE exposed groups (non-combined) were injected with the same volume of DMSO, and the injected Pacific oysters were incubated in the original glass for 24 h. The rearing conditions were the same as the pre-domestication conditions, and samples were collected for subsequent experiments.

#### Real-time quantitative PCR analysis of expression of E3 ubiquitin ligase and the NF-κB signalling pathway- and autophagy related-genes in Pacific oysters after PHE and combined PHE and CAPE exposures

After 24 h of exposure, six oysters were selected each from the DMSO control, PHE exposure, and combined PHE and CAPE exposure groups, and 800 μL of haemolymph was collected from the blood sinus of each oyster using a 1 mL sterile syringe. The samples were mixed into a 10 mL centrifuge tube, and the cell concentration was adjusted to 1 × 10^6^ with PBS (pH7.4). After centrifugation at 860 × g for 5 min, the supernatant was discarded. The cells were isolated using a TRIzol reagent (Invitrogen) and further purified using a Total RNA Isolation kit (TaKaRa, Shiga, Japan), following the manufacturer’s protocol with minor modifications. Briefly, the cells were mechanically disrupted in the presence of 1 mL of TRIzol using grinding rods. RNA was eluted in 35 μL of RNase-free water (TaKaRa) and stored at −80°C. Residual genomic DNA contamination was removed during RNA clean-up using RNase-free DNase I digestion, as instructed by the manufacturer (TaKaRa). RNA concentration and purity were then measured using a spectrophotometer (NanoDrop ND-2000c; Thermo Fisher Scientific, Waltham, MA, USA) and 1% agarose gel electrophoresis. Only high-purity samples (OD 260/280 > 1.8, OD 260/230 > 1.8) were subsequently processed. Equal amounts of RNA (1 μg) were used to synthesise cDNA using the PrimeScript™ RT kit (TaKaRa). Aliquots of the RT mixture were diluted to 1/10 with nuclease-free water (Sigma) before use. The resulting undiluted and diluted cDNA was stored at −20°C.

Based on the gene sequences for the Pacific oyster in GenBank, primers for the target genes were designed using Primer Premier 5.0 software (PREMIER Biosoft, Palo Alto, CA, USA; Supplementary Data 1). Six internal reference genes were selected and tested for transcriptional stability, and 40S_s3 and 40S_s9 (M value = 0.680 and 0.708, respectively) were selected as the internal reference genes using geNorm analysis (Wang et al., 2020).

The relative gene transcript levels in the lymphocytes from the DMSO control and exposed oysters were investigated using qRT-PCR in Eppendorf MasterCycler® ep RealPlex4 (Eppendorf, Wesseling- Berzdorf, Germany), with primers specific to the *C. gigas* genes. qPCR was performed using the SYBR Green Mix Kit (TaKaRa, Dalian, China). The qPCR reaction system was 20 μL, containing 0.8 μL of the forward and reverse primers (0.8 μM), 2 μL cDNA samples, 0.4 μL ROX reference dye, 10 μL 2 × SYBR Premix Ex Taq (Takara Bio, Shiga, Japan), and 6.0 μL nuclease-free water. PCR amplification was performed using the following cycling program: 30 s at 95°C, 40 cycles of 5 s at 95°C, 30 s at 60°C, and 1 min at 60°C, as instructed by the manufacturer. Melting curve analysis was performed on each PCR product to determine the specificity of the amplified fragments. The expression level of each target gene was normalised to the geometric mean of the mRNA contents of the two internal reference genes using the 2^-ΔΔCT^ method (Livak et al., 2001).

### Quantitative real-time PCR analysis of expression of E3 ubiquitin ligases and NF-κB signalling pathway- and autophagy-related genes in murine macrophage RAW264.7 cells exposed to PHE

To verify the regulatory mechanism of PHE-induced autophagy in the lymphocytes, quantitative real-time PCR was performed to analyse the expression of E3 ubiquitin ligases and the NF-κB signalling pathway- and autophagy-related genes in murine macrophage RAW264.7 cells exposed to PHE. Cells were cultured in 25 T culture flasks with 3 mL DMEM medium (Invitrogen, 111960-044), and there were 10, 50, and 250 μg/L PHE exposure groups and a DMSO control group, with incubation at 37°C and 5% CO_2_ for 24 h. After the cells were transferred to five 25 T culture flasks with 1 mL trypsin, the culture medium was discarded after half an hour, and the cells were transferred to a 1.5 mL centrifuge tube after mixing with 1 mL TRIzol (Invitrogen). Subsequent gene expression detection experiments were performed as described in Section “Real-time quantitative PCR analysis of expression of E3 ubiquitin ligase and the NF-κB signalling pathway- and autophagy related-genes in Pacific oysters after PHE and combined PHE and CAPE exposures”.

### Effect of PHE exposure on the total number of lymphocytes, apoptosis rate, phagocytosis, and ROS level

Eight oysters from each group after exposure to the PHE for 24 h and 48 h were selected, and 800 μL of haemolymph was extracted from the blood sinus using a 1 mL sterile syringe. The samples from each group were mixed in a 10 mL centrifuge tube with 600 μL of anticoagulant (PBS containing 10 mg/mL sodium heparin), and each sample was divided into four portions to monitor the number of total haemolymph cells, phagocytic capacity, apoptosis rate, and ROS content.

#### Total number of lymphocytes

The prepared samples of lymphocytes (300 µL) that were extracted from each group were fixed in 10 µL of paraformaldehyde solution (Sigma-Aldrich, G5882) for 10 min at 4°C. Then, 10 µL of the fixed lymphocytes were added dropwise to the haemocytometer plate, the number of haemocytes was counted using microscopic observations at 60 × magnification, the total number of lymphocytes in each group of samples was counted, and the measurements were repeated at least three times. The total number of haemocytes was calculated as follows: 5 middle compartment haemocytes × 25/5 × 10^4^ × dilution times (pcs/mL).

#### Apoptosis rate of the lymphocytes

After centrifugation of the lymphocytes at 800 × g for 10 min at 4°C, the supernatants were removed, and the cell precipitates collected. The cells were gently resuspended in PBS three times. After incubation with 195 µL of Annexin V-FITC conjugate, 5 µL of Annexin V-FITC and 10 µL of propidium iodide were added for staining. The sample was mixed gently and incubated for 10–20 min at 20–25°C away from the light and then was placed in an ice bath prior to measurement. The green fluorescence (FL1) for Annexin V-FITC and the red fluorescence (FL3) for propidium iodide (PI) were detected using flow cytometry, with the parameters identified by Picot et al. (Picot et al., 2019).

#### Phagocytic capacity of the lymphocytes

Microbial labelling was performed using fluorescein isothiocyanate (FITC, Aladdin, Shanghai, China), and 100 µL of 1 × 10^9^ CFU/mL *Staphylococcus aureus* was fixed with 20 µL of formaldehyde for 10 min, followed by centrifugation at 2380 × g for 5 min, and then the supernatants were removed. Bacterial precipitates were resuspended with FITC working solution (concentration 1 mg/mL, prepared with 0.1 M NaHCO_3_ solution, pH9.0). After incubation for 3 h at room temperature with shaking, the samples were washed three times with PBS, the bacterial concentration was adjusted to 1 × 10^8^ CFU/mL, and the samples were stored at 4 ℃ prior to the experiments.

The lymphocytes were washed three times with PBS, adjusted to a concentration of 1 × 10^6^ cells/mL with PBS, and incubated with an equal volume of FITC-labelled bacteria for 1 h at room temperature with light shaking. After centrifugation at 800 × g for 10 min at 4 ℃, the cells were washed three times with PBS, then resuspended in PBS, and filtered through a strainer to a tube. The phagocytic capacity of the lymphocytes was determined using flow cytometry with the same parameters as described in Section “*Apoptosis rate of the hemocytes*”. The phagocytic capacity of the lymphocytes included the phagocytic rate and phagocytic index. The lymphocyte phagocytic rate (PR) is the ratio of the number of phagocytes to the total number of lymphocytes. The phagocytic index is the average fluorescence value of the labelled microorganisms in the phagocytes.

#### ROS levels in the lymphocytes

The samples were washed three times with PBS and finally adjusted to a concentration of 1 × 10^6^ cells/mL with PBS, followed by co-incubation with the fluorescent probe DCFH-DA from the assay kit (Beyotime, S0033S, Shanghai, China). ROS levels were detected using a flow cytometer (Beckman FC500-MPL, USA), according to the method of Zhao et al. (Zhao et al., 2019).

### Statistical analysis

All data are expressed as the mean ± standard deviation (SD). Statistical analysis was performed using SPSS software (Chicago, IL, USA), and the data were tested for homogeneity of variance using Levene’s test. Statistical differences between the exposed and control groups were tested using one-way ANOVA with LSD (one-way ANOVA), with 0.01 < P < 0.05, indicating a significant difference (marked with *) and P < 0.01, indicating a highly significant difference (marked with **).

### Resource availability

#### Lead contact

Further information and reasonable requests for resources and reagents should be directed to and will be fulfilled by the lead contact, Pengfei Cui.

#### Materials availability

Requests for materials should be made via the lead contact. All unique/stable reagents generated in this study are available from the lead contact without restriction.

### Data and code availability

- ***Data***: All data reported in this paper will be shared by the lead contact [Pengfei Cui] upon reasonable requests.
- ***Code***: This paper does not report original code.
- ***Additional Information***: Any additional information required to reanalyze the data reported in this article is available from the lead contact upon request.

## Acknowledgements

This work was supported by the national key research and development program of China (2017YFC1600705). The authors thank Dr. Ming Gao of University of Soochow, for his technical aids in HPLC. The authors also thank Ocean University of China for providing funds for highly talented young researchers.

## Author contributions

Conceptualization, P.C., Z.Z. and S.R.; Methodology, P.C., W.W. and R.T.; Formal Analysis, Z.G., Z.Z., H.M., H.D., W.W. and R.T.; Investigation, Z.G., Z.Z., H.M., H.D., P.C., R.T. and W.W.; Data Curation, Z.G., Z.Z., H.M., H.D. and P.C.; Visualization, Z.G. and H.M.; Resources, P.C. and S.R.; Writing – Original Draft, Z.G., H.M. and P.C.; Writing – Review & Editing, Z.G., H.M. and P.C.; Funding Acquisition, P.C. and S.R.; Supervision, P.C. and S.R.

## Abbreviations

PAHs: Polycyclic aromatic hydrocarbons
POPs: persistent organic pollutant
AMPK: AMP-activated protein kinase
PI3K: phosphatidylinositol 3-kinase
KK: IκB kinase
κB: NF-κB inhibitor
UPS: ubiquitin-proteasome system
ALP: autophagy-lysosomal pathway
CLSM: confocal laser scanning microscopy
GO: Gene Ontology
THC: total number of blood lymphocyte
C. gigas: Pacific oysters
DMEM: Dulbecco’s modified Eagle’s medium
PBS: phosphate-buffered saline
CAPE: caffeic acid phenyl ether
FITC: fluorescein isothiocyanate
PR: phagocytic rate.

