## Supplemental Figures and Tables for "Cytoprotective roles of “E3 ubiquitin ligases-NF-κB-autophagy” axis in Pacific oysters *Crassostrea gigas* exposed to phenanthrene"

**Supplementary Table 1.** The sequences of primers used for real-time PCR validation

| Gene | GenBank ID | Primer Sequence (5'-3') |
|---|---|---|
| *40S_s3-like* | 105335873 | F: CCTGTTATGGAGTGCTACGGTTTA TC |
| | | R: CATTGACTTGGCTCTCTGTCCTC |
| *40S_s9-like* | 105341101 | F: CCTGTTCCCTTCTTGGCATTCTT |
| | | R: TGACTTCTCCCT AGATCACCATAC |
| *CgNF-κB* | 111125961 | F: GAAGGCAAAGGGAGGTGATGAG |
| | | R: GGTGTGCGGAAGACAATGGC |
| *CgIκκ-α* | 105348304 | F: TCTCACACCCACACACCTATGC |
| | | R: AGTAGTTTTCCACCAGGGGATAAG |
| *CgLC3* | 59371002 | F: ATGGCGGCAGGTTCAATTTCGGA |
| | | R: GCTGACACAAAGTTTTCCATACTCCAGAGGA |
| *CgSQSTM1* | 105346676 | F: AGGGAATGAGAAGGCCGAAA |
| | | R: CCTCAAGCAACTCCTCTCCA |
| *CgHUWE1* | 105318881 | F: GACAACACAACCACAGTGCC |
| | | R: TCCCCGTTCTCCATCCTCAT |
| *CgTRIM36* | 105342581 | F: TGTTCCTAATGTGAGGGTCCAC |
| | | R: TTCACGCGCAGATGTATGGT |
| *M-β-actin* | 11461 | F: GAGACCTTCAACACCCCAGC |
| | | R: ATGTCACGCACGATTTCCC |
| *M-gapdh* | 14433 | F: AGGTCGGTGTGAACGGATTTG |
| | | R: GGGGTCGTTGATGGCAACA |
| *M-NF-κB* | 18033 | F: AATCAAGAGCTCCGAGACGC |
| | | R: GGCATTTCTTCAGTTGCTCCA |
| *M-Iκκ-α* | 16150 | F: GGCACCCAATGATTTGCCAC |
| | | R: CTGGCGCCGATGTCACTC |

| | | |
|---|---|---|
| *M-LC3* | 67443 | F: ACAAGGGAAGTGATCGTCGC<br>R: TCGCTCTATAATCACTGGGATCT |
| *M-SQSTM1* | 18412 | F: ACCAAGATCCCAGTGATTATAGAGC<br>R: TGCAAGCGCCGTCTGATTAT |
| *M-HUWE1* | 59026 | F: GAGGTTCTCCTGGGATCACG<br>R: GCTACTGAAGGACCGGCTAAG |
| *M-TRIM36* | 28105 | F: TGGCGTCAGACAGCTCAAAT<br>R: ATTACGCTTCCAGCCTGGTC |

F: Forward primer sequence, R: Reverse primer sequence

**Supplementary Table 2.** Partial difference protein between DMSO group and 50 μg/L exposed group

| Difference Protein | Pvalue | Up/Down |
|---|---|---|
| E3 ubiquitin-protein ligase HUWE1-like isoform X1(XP_011414495.1) | 0.0011260<br>51 | up |
| 4-coumarate--CoA ligase-like 7(XP_011422712.1) | 0.0032967<br>82 | up |
| piwi-like protein 1 isoform X2(XP_019926528.1) | 0.0042358<br>15 | up |
| copper transport protein ctr6(XP_011412212.1) | 0.0049322<br>63 | down |
| fatty-acid amide hydrolase 2-A(XP_011454525.1) | 0.0061607<br>37 | up |
| 60 kDa heat shock protein (XP_011456445.1) | 0.0068379<br>48 | up |

| | | |
|---|---|---|
| glycine dehydrogenase (XP_019925034.1) | 0.007241992 | up |
| serine/threonine-protein kinase 31(XP_011430517.1) | 0.007587151 | up |
| E3 ubiquitin-protein ligase TRIM36(XP_011438896.1) | 0.010055006 | up |
| mitochondrial 2-oxodicarboxylate (XP_011413362.1) | 0.010694475 | up |
| centrosomal protein POC5(XP_019919480.1) | 0.012456936 | down |
| nuclear receptor coactivator 4(XP_011431072.1) | 0.0272 | up |
| mitochondrial amidoxime reducing (XP_011439675.1) | 0.0272 | up |
| ATP-binding cassette sub-family D(XP_011414032.1) | 0.015751149 | up |
| annexin A7(XP_011442013.1) | 0.019598596 | up |
| Catalase (XP_011443987.1) | 0.021959654 | up |
| methionine adenosyltransferase 2(XP_011432001.1) | 0.02251104 | down |
| ubiquitin-like modifier-activating enzyme ATG7(XP_019927770.1) | 0.047345374 | up |

**Supplementary Figure 1.** Concentration of PHE in each tissue of *Crassostrea gigas* and its hemolymph fluid: (A) Concentration of PHE in the muscle (μg/kg) after 24 h and 7 d exposure, (B) Concentration of PHE in the coat membrane (μg/kg) after 24 h and 7 d exposure, (C) Concentration of

PHE in the visceral mass (μg/kg) after 24 h and 7 d exposure, (D) Concentration of PHE concentrations in gills (μg/kg) after 24 h and 7 d exposure,(n=3).

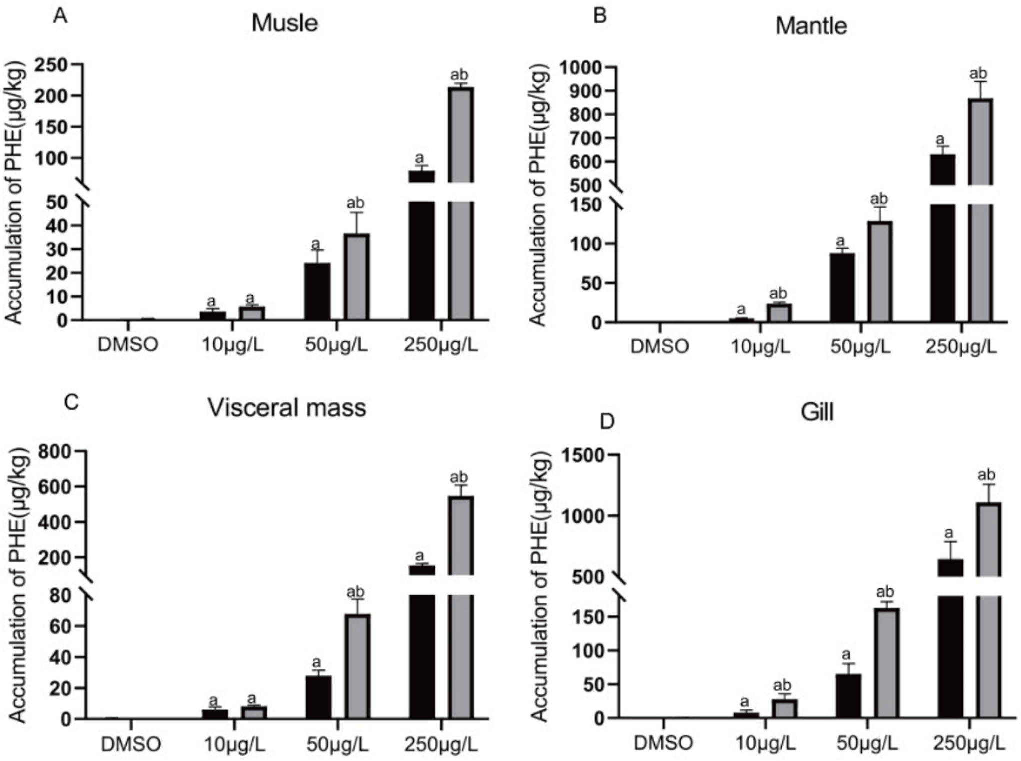


**Supplementary Figure 2.** CAT and SOD enzyme activity indices in Pacific oysters, (A) CAT enzyme activity indices of visceral masses after 24h, 48h, and 72h exposure (B) CAT enzyme activity indices of gills, visceral masses, and mantle after 24h, 48h, and 72h exposure, (n=6).

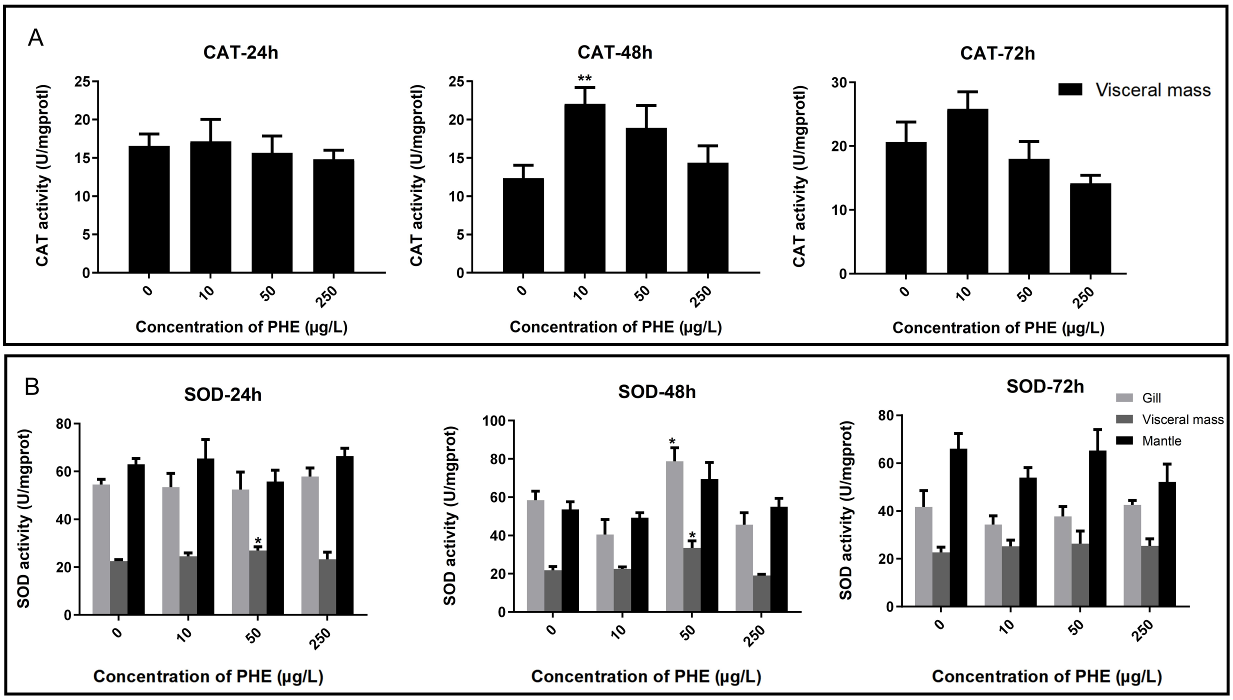
A
CAT-24h
CAT-48h
CAT-72h
CAT activity (U/mgprotl)
Visceral mass
**
0
10
50
250
Concentration of PHE (μg/L)
B
SOD-24h
SOD-48h
SOD-72h
SOD activity (U/mgprot)
Gill
Visceral mass
Mantle
*
Concentration of PHE (μg/L)
